# Linkage: an interactive shiny app and R package for linking of DNA regulatory peaks to genes

**DOI:** 10.1101/2024.04.24.590756

**Authors:** Zenghui Liu, Shaodong Chen, Tianting Li, Chao Zhang, Yuyan Luo, Junxi Zheng, Zixiao Lu, Jin Yang, Siwen Xu

## Abstract

**Summary:** Existing studies have demonstrated that the integration analysis of transcriptomic and epigenomic data can be used to better understand the onset and progression of many diseases, as well as identify new diagnostic and prognostic biomarkers. However, such investigations on large-scale sequencing data remain challenging for researchers or clinicians with limited bioinformatics knowledge. To facilitate the interpretation of gene regulatory landscape, we developed an R Shiny application and R package [**Link**ing of **a**tac-seq to **g**ene **e**xpression data (Linkage)] for exploring and visualizing potential cis-regulatory elements (CREs) of genes based on ATAC-seq and RNA-seq data. Linkage offers six modules to systematically identify, annotate, and interpret potential gene regulatory elements from the whole genome step by step. Linkage can provide interactive visualization for the correlation between chromatin accessibility and gene expression. More than that, Linkage identifies transcription factors (TFs) that potentially drive the chromatin changes through identifying TF binding motifs within the CREs and constructing trans-regulatory networks of the target gene set. This powerful tool enables researchers to conduct extensive multiomics integration analysis and generate visually appealing visualizations that effectively highlight the relationship between genes and corresponding regulatory elements. With Linkage, users can obtain publishable results and gain deeper insights into the gene regulatory landscape.

**Availability and implementation:** ‘Linkage’ is freely available as a Shiny web application (https://xulabgdpu.org.cn/linkage) and an R package (https://github.com/XuLab-GDPU/Linkage). The documentation is available at (https://aicplane.github.io/Linkage-tutorial/).

## 1. Introduction

With the rapid advancement of high-throughput sequencing technologies and bioinformatics analysis, there is a growing awareness of the crucial role played by the genomic non-coding regions in gene regulation (Dong, et al., 2023). Vast number of gene regulatory elements, such as transcription factors binding sites (TFBS), are enriched in these regions. Chromatin accessibility represents the degree of physical contact of gene regulatory elements with chromatin DNA and is widely studied in the epigenetic research (Domcke, et al., 2020). Only active cis-regulatory elements (CREs) can be accessible by the trans-regulatory factors such as transcription factors (TFs). Assay for transposase-accessible chromatin with high-throughput sequencing (ATAC-seq) is a high-throughput technology that employs an engineered Tn5 transposase to map and quantify genome-wide chromatin accessibility (Buenrostro, et al., 2015). Higher chromatin accessibility means more active DNA-protein binding events that happen at these regions for bulk ATAC-seq data analysis. On the other hand, bulk mRNA sequencing (RNA-seq) is the most common application to estimate and quantify gene expression level, which has broad utilities in differential expression analysis, cancer classification, disease diagnosis, and optimizing therapeutic treatment (Alharbi and Vakanski, 2023; Buzdin, et al., 2020).

To date, many studies have characterized gene regulatory landscapes by integrating ATAC-seq and RNA-seq data (Li, et al., 2024; Xu, et al., 2020). However, this kind of work remains very challenging for scientists that without enough experience in high-throughput sequencing data analysis. The existing web tools only support analyzing datasets that from public domains such as The Cancer Genome Atlas (TCGA) and The Encyclopedia of DNA Elements (ENCODE) (Abascal, et al., 2020; Wu, et al., 2020).

Here, we introduce Linkage, a user-friendly interactive R Shiny web-based application for exploring and visualizing potential gene regulatory elements based on ATAC-seq and RNA-seq data. Linkage allows users to upload customized data or re-analyze public datasets. The main feature for Linkage is to identify potential gene regulatory elements for the whole genome by performing correlation analysis between chromatin accessibility and gene expression from the same sample. Additional modules are developed to allow users to perform deeper and more systematic analysis for the links between ATAC-seq peaks and target genes. In brief, Linkage provides an integrative analysis of ATAC-seq and RNA-seq data for potential gene regulatory elements identification. Analysis and visualization results are returned to the web page and can be downloaded in PDF, PNG, JPEG, CSV, and TXT formats. The workflow and typical output schema are shown in Fig. 1.

**Fig. 1.**
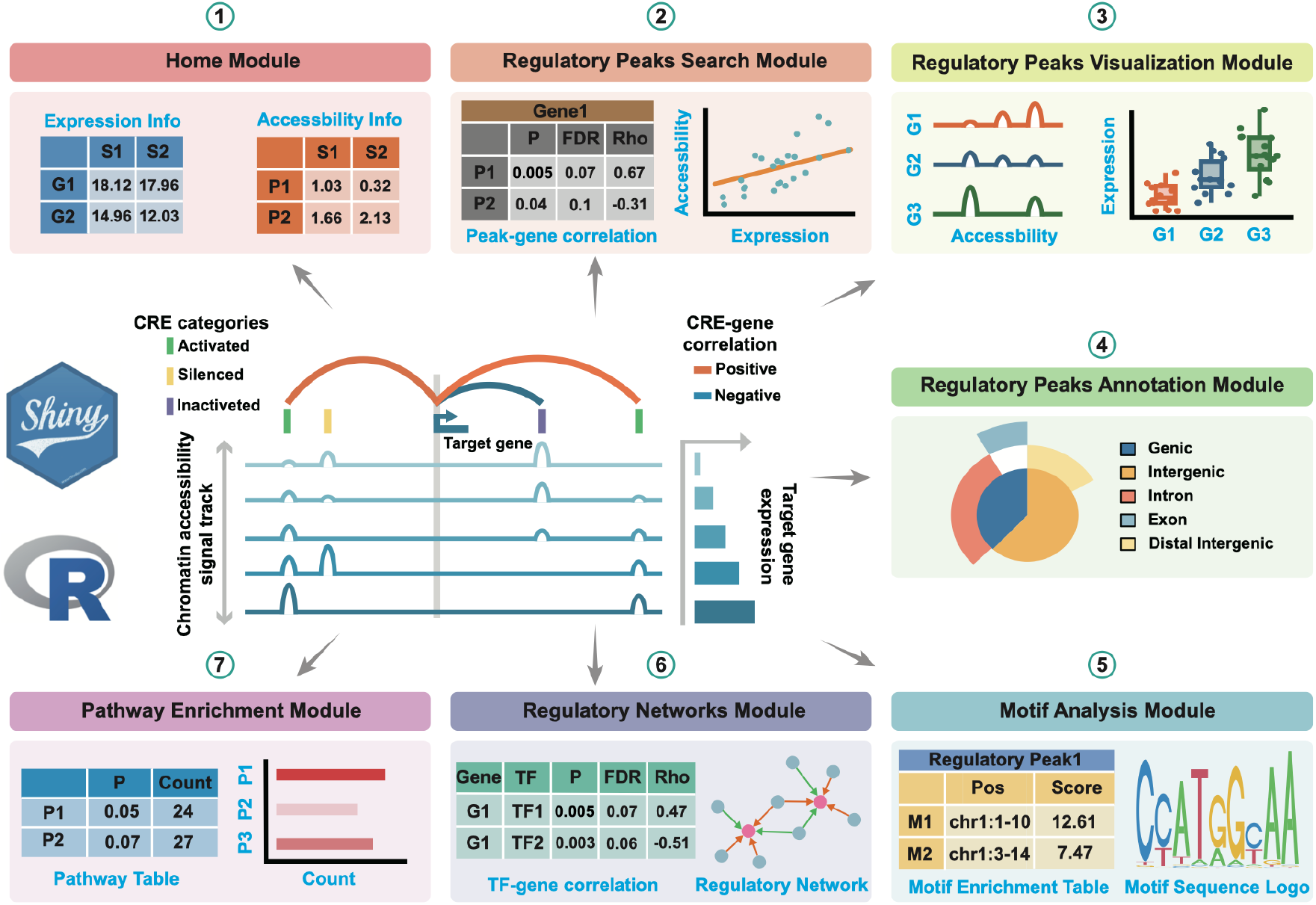
Overview of functionality and workflows with Linkage.

## 2. Modules

### Home Module

The Home Module contains the brief instruction and Import Data panel of Linkage. Linkage provides two public multi-omics datasets, i.e., the ***TCGA Breast Cancer cohort*** dataset and the ***GSE121589*** dataset, for users to re-analyze them. The ***TCGA Breast Cancer cohort*** is a human dataset that contains bulk RNA-Seq and ATAC-Seq data from 72 patients with breast cancer (Corces, et al., 2018). The ***GSE121589*** is a mouse dataset that constitutes bulk RNA-Seq and ATAC-Seq data from 33 mice Muscle Stem Cells (MuSCs) (Shcherbina, et al., 2020). Apart from the provided public datasets, Linkage also enables users to upload and explore their own data.

Only two tab-delimited text/csv input files (a chromatin accessibility matrix and a gene expression matrix) are required before running Linkage. The gene expression matrix file is a tab-delimited multi-column data matrix, where the first column represents gene symbols and the following columns represent normalized or raw expression levels of genes for each sample. The chromatin accessibility matrix file is also a tab-delimited multi-column data matrix, where the first three columns represent chromosome name, start and end coordinates on the chromosome of the peaks respectively, and the remaining columns represent normalized or raw chromatin accessibility levels of peaks for each sample. Each element of ready-to-analysis files will appear in Chromatin Accessibility Matrix panel and Gene Expression Matrix panel.

### Regulatory Peaks Search Module

The Regulatory Peaks Search Module is the key module of Linkage application that allows users to detect all potential regulatory DNA regions for a specific gene (Fig. 1). Linkage adopted the same procedure as in TCGA (Corces, et al., 2018) to associate regulatory regions with the predicted target genes. Specifically, given the input gene and targeting DNA regions, Linkage will automatically perform canonical correlation analysis between each quantitative chromatin accessibility measure in the region and the quantitative expression level of the gene across all samples. Users can easily adjust the regions of interest and customize the correlation algorithm (Spearman / Pearson / Kendall). Then, all the statistically significant results are listed in the Potential Cis-regulatory Regions panel. With clicking on a specific entry of this panel, users can view the scatter plot of quantitative chromatin accessibility and gene expression from the Correlation Plot panel. The corresponding rho and FDR for the correlation analysis will also be shown on the scatter plot. A positive correlation indicates that more accessible of open chromatin regions appear to have higher expression level of genes. Such finding suggests that the CREs in such regions could potentially activate the expression of target genes. On the contrary, a clear negative correlation suggests that the target gene associated with the chromatin accessibility regions might be repressed by the CREs in these regions.

### Regulatory Peaks Visualization Module

The Regulatory Peaks Visualization Module allows users to visualize the coverage of mapped ATAC-seq reads around a given specific regulatory peak, as well as the corresponding quantitative expression of the target gene of this regulatory peak. Users initially select a regulatory peak that is obtained from the Regulatory Peaks Detection Module. Linkage then categorizes samples into five groups based on the quantitative chromatin accessibility of the specific regulatory peak, ranging from low to high for each individual sample. The coverage track of mapped ATAC-seq reads and the expression boxplot of the target gene for each group will be shown simultaneously.

### Regulatory Peaks Annotation Module

The Regulatory Peaks Annotation Module allows users to visualize the annotation of the predicational regulatory peaks for genes that are given in the previous module. Once users click ‘Annotating Regulatory Peaks’, Linkage will perform the annotation of all predicational regulatory peaks in terms of genomic location features, including whether the peaks are in the TSS, Exon, 5’ UTR, 3’ UTR, Intronic or Intergenic and the position and strand information of the nearest gene of the peaks. To effectively visualize the overlaps and distribution in the annotation for peaks, Linkage also produces the upsetplot that is adopted from the ChIPseeker package (Wang, et al., 2022).

### Cis-Regulatory Elements Detection Module

After identifying potential regulatory peaks, researchers can further explore which specific CREs are activated with Linkage. CREs and the TFs that binding on them play a central role in gene transcription regulation, which can be detected by ATAC-seq data. The Cis-Regulatory Elements Detection Module supports users to visualize the enriched TFBS within potential regulatory peaks. With clicking on a specific regulatory peak, users can view the location and binding score information of each enriched TFBS of this DNA region from the Motif Enrichment Table. Once users select one TFBS of this table, the corresponding sequence logo of this CRE will appear in the Sequence-logo Plot panel.

### Gene Regulatory Network Module

The interplay between CREs and genes generates complex regulatory circuits that can be represented as gene regulatory networks (GRNs). Studies of GRNs are crucial to understand how cellular identity is established, maintained and disrupted in disease. The Gene Regulatory Network Module helps users to visualize GRNs that nodes are genes and corresponding CREs, which inferred from previous analysis of Linkage. First, users can input a list of interested genes or upload a gene-list file which was obtained from the previous analysis. Then users can adjust a series of parameters in association with building the GRN, including types of gene symbols, calculation methods and thresholds of interactions between the nodes (edges of the GRNs). Once users click the ‘Build GRNs’ button, Linkage will perform canonical correlation analysis of quantitative expression level between each interested gene and their potential CREs. The significant calculation results of correlation analysis are shown in the Gene-TF Table panel. Meanwhile, Linkage produces the corresponding informatic and interactive GRNs that are adopted from the visNetwork package (Almende, et al., 2019). Users can further easily change network layouts, select subnetworks, and save the GRNs as spreadsheets with interaction score or plots. Furthermore, Linkage allows users to upload the previously saved Gene-TF Table file for rebuilding the GRN, which is very time-consuming.

### Pathway Enrichment Analysis Module

The Pathway Enrichment Analysis Module supports users to visualize tabular and graphical pathway enrichment results of interested gene lists that are previously produced from other modules of Linkage. The pathway enrichment analysis can link these genes with underlying molecular pathways and functional categories such as gene ontology (GO) (Consortium, 2019) and Kyoto Encyclopedia of Genes and Genomes (KEGG) (Kanehisa and Goto, 2000). Within this module, users can input a list of interested genes and set four key parameters (i.e., adjusted pvalue cutoff, qvalue cutoff, minimal size of annotated genes for testing, and maximal size of annotated genes for testing) for pathway enrichment analysis following the implementation of the clusterProfiler package (Yu, et al., 2012). Then once users click the ‘Build Pathway Enrichment Analysis’ button, Linkage automatically performs GO and KEGG enrichment analysis. The corresponding enrichment categories will be returned in the GO/KEGG Enrichment Table. To interpret the functional results from multiple perspectives, Linkage implements several different visualization methods that obtained from the GO/KEGG Enrichment Table, which adopting from the enrichplot package (Wu, et al., 2021)

## 3. Case Study

To better show the functionalities of Linkage, we collect multi-omics profiles of human breast cancer and mice MuSCs for experiments. For the human breast cancer dataset, we collect 72 samples with matched chromatin accessibility and gene expression data. For each detected expressed gene, we asked which regulatory regions might have contributed to the expression of this gene in breast cancer patients. To address this question, we implemented canonical correlation analysis between the quantitative chromatin accessibility measure and the gene expression level across all samples. The Spearman correlation coefficient, r, was defaultly used to evaluate whether the potential regulatory regions were significantly correlated with the gene expression. A total of 60,483 expressed genes were analyzed. All peaks whose summit were located within 500 kbp from a gene’s TSS were considered by default. A conservative FDR cutoff of 0.01 was used to avoid false positives. This analysis identified a total of 489,132 regulatory regions for 18,104 genes and a complete list of these regulatory regions can be found in Additional file 1: Table S1. We also collect matched chromatin accessibility and gene expression data of 33 mice MuSCs. A total of 1475 regulatory regions for 910 genes are identified with the same protocol and a complete list of these regulatory regions can be found in Additional file 2: Table S2. Taken together, these results demonstrate that the Linkage application can identify gene expression-associated regulatory regions from multi-omics data, which could potentially implicate regulatory elements responsible for cancer development.

## 4. Conclusions

Linkage is a web-based Shiny application/R package that can utilize large-scale multi-omics datasets for identification and prioritization of DNA regulatory elements from whole genomic regions. Linkage helps users to perform canonical correlation analysis between ATAC-seq accessibility and gene expression across all samples for regulatory peaks identification. The visualization module and annotation module provide user-friendly graphical interfaces to visualize the ATAC-seq genome tracks and genomics location features of a given regulatory peak. The motif module helps assess the probability of potential DNA regulatory elements that can exist within individual regulatory peak. Linkage web application also supports users to explore the GRNs and molecular pathways composing from DNA regulatory elements and target genes that were collected by the previous analyses. Thus, Linkage web application is ideal for integrative analysis when users have limited bioinformatics knowledge. In addition, we provide more customizable parameters within the Linkage package to ensure the analysis efficiency for those with bioinformatics skills. Furthermore, the R package implementation of Linkage enables users to easily integrate our algorithm into existing analytical pipelines within the same R environment with flexible graphing options. A more comprehensive tutorial for running the web application and R package of Linkage is available either as webserver (https://aicplane.github.io/Linkage-tutorial/) or GitHub (https://github.com/xulabgdpu/linkage). In conclusion, with the increasing popularity of higher-resolution sequencing strategies such as single-cell technologies, integrative multi-omics data analysis will undoubtedly receive more and more attention. We will keep on developing and upgrading the Linkage network resource to benefit the omics data integration research community.

## Notes

https://xenabrowser.net/datapages/?cohort=GDC%20TCGA%20Breast%20Cancer%20(BRCA)&removeHub=https%3A%2F%2Fxena.treehouse.gi.ucsc.edu%3A443

https://www.ncbi.nlm.nih.gov/geo/query/acc.cgi?acc=GSE121589

### Competing Interest Statement

The authors have declared no competing interest.

